# Electron-density informed effective and reliable *de novo* molecular design and lead optimization with ED2Mol

**DOI:** 10.1101/2024.12.18.629081

**Authors:** Mingyu Li, Kun Song, Mingzhu Zhao, Gengshu You, Jie Zhong, Mengxi Zhao, Arong Li, Yu Chen, Guobin Li, Ying Kong, Jiacheng Wei, Zhaofu Wang, Jiamin Zhou, Hongbing Yang, Shichao Ma, Hailong Zhang, Irakoze Loïca Mélita, Weidong Lin, Yuhang Lu, Zhengtian Yu, Xun Lu, Jixiao He, Yujun Zhao, Jian Zhang

## Abstract

Generative drug design opens new avenues for discovering novel compounds within the vast chemical space rather than conventional screening against limited compound libraries. However, the practical utility of the generated molecules is frequently constrained, as many designs prioritize a narrow range of pharmacological properties while neglecting physical reliability, which hinders the success rate of subsequent wet-lab evaluations. To address this, we propose ED2Mol, a deep learning-based approach that leverages fundamental electron density information to improve *de novo* molecular generation and lead optimization. The extensive evaluations across multiple benchmarks demonstrate that ED2Mol surpasses existing methods in terms of generation success rate and >97% physical reliability. It also facilitates automated lead optimization that is not fully implemented by other methods using fragment-based strategies. Furthermore, ED2Mol exhibits generalizability to more challenging, unseen allosteric pocket benchmarks, attaining consistent performance in both *de novo* molecule generation and lead optimization. More importantly, ED2Mol has been applied to various real-world essential targets, successfully identifying wet-lab validated bioactive compounds, ranging from FGFR3 orthosteric inhibitors to CDC42 allosteric inhibitors and GCK allosteric activators. The directly generated binding modes of these compounds with target proteins are close to predictions through molecular docking and further validated via the X-ray co-crystal structure. All these results highlight ED2Mol’s potential as a useful tool in realistic drug design with enhanced effectiveness, physical reliability, and practical applicability.

## Main

Innovative drug discovery has long been recognized as a time-intensive, costly, and high-risk endeavor^1,2^. In general, this process is categorized into four stages: hit identification, lead optimization, preclinical evaluation, and clinical trials. Among these, hit/lead compound discovery serves as a cornerstone, as identifying a highly potent active compound can substantially enhance the effectiveness of subsequent clinical evaluation/trials^3^. Over the past several decades, the majority of active compounds have been identified through high-throughput screening against established compound libraries, which typically consist of between 10^4^ and 10^10^ molecules^4-6^. Nowadays, after extensive and prolonged screening of such limited-diversity libraries through various campaigns, discovering active compounds with novel scaffolds has become increasingly challenging.

Alternatively, *de novo* molecular design—where active compounds are generated from scratch—offers a promising strategy to explore a much broader chemical space (∼10^60^) beyond existing libraries^7,8^. With the explosion of available protein structures and advancements in deep generative models (DGMs), structure-based molecular design, particularly in the realm of structure-aware *de novo* molecular design, has been significantly empowered^9^. Given that active compounds tightly bind to specific binding pockets within protein structures, structure-aware DGMs are considered more efficient for molecular generation than commonly used structure-unaware, ligand-based DGMs^10^. To some extent, structure-based DGMs have indeed shown success in producing ligands that potentially bind more tightly, with many exhibiting better computational binding scores than reference experimental ligands (generation success rate). However, a closer examination of the ligands generated by existing models reveals several challenges, outlined as follows, with further discussion provided later: (1) DGMs primarily rely on high-quality big datasets, yet the limited dataset of experimental protein-ligand complex hinders the training of robust DGMs applicable to diverse binding pockets. (2) In terms of intermolecular reliability, many generated molecules present binding poses that clearly clash with the receptor and/or undergo drastic reorientation upon re-docking, indicating that DGMs fail to capture optimal interatomic interactions, and the directly generated poses often represent sub-optimal binding modes^11^. (3) In terms of intramolecular reliability, a notable proportion of generated molecules display physiochemically implausible topologies and/or complex substructures, probably leading to poor synthetic accessibility and/or high toxicity, thus posing a considerable failure risk in subsequent preclinical evaluations^12^. (4) The generated compounds often act as hit compounds, exhibiting moderate potency with potential for further lead optimization. Unfortunately, most current *de novo* design models lack implementations of lead optimization networks. (5) Although most DGMs have been computationally validated to produce ligands with excellent affinity, they struggle to accurately demonstrate the bioactivity and optimal binding modes of generated ligands without wet-lab experiments, thereby restricting their practical application in the real-world drug discovery pipelines.

Integrating domain-specific insights into DGMs is increasingly recognized as a promising approach to address their robustness challenges^13,14^. Structural biologists contribute to this effort by elucidating optimal interatomic interactions through the determination of protein-ligand complex structures, particularly the ligand electron density (ED) map. From a microscopic perspective, the ED encapsulates the fundamental mechanisms governing protein-ligand interaction forces, such as the formation of Van der Waals interactions and hydrogen bonds, which arises from the “pull-push” effects of electrons between atoms^15,16^. From a macroscopic perspective, the ED cloud outlines the spatial complementarity between the ligand and the receptor, as conceptualized by the well-known “lock-and-key” theory of ligand binding^17,18^. Consequently, incorporating ED into DGMs would not only enable a more generalized and accurate modeling of interatomic interactions via unified electromagnetic forces, but also foster a deeper understanding of the global complementary environment of the binding pocket. Unlike existing models that determine locally optimal atoms or motifs by considering only neighboring pocket atoms, this strategy might reduce the risk of sub-optimal binding modes. Once a ligand’s ED is obtained, structural biologists fit its three-dimensional structure into the ED map. One commonly used automated tool for this purpose is LigandFit, embedded in Phenix^19,20^. LigandFit initially decomposes the ligand into fragments, then initially places the most rigid part of the ligand (namely the core) and iteratively attaches the remaining fragments to the core following the ED (Fig. 1a). Such fragment-based strategy is also prevalent in drug design (FBDD), as they provide a more efficient means to explore a broad and synthetically accessible molecular chemical space by assembling synthesizable building fragments based on reasonable assembly rules^21^. This strategy has the potential to resolve the aforementioned intramolecular reliability issues. Moreover, it can be seamlessly applied in lead optimization, where fragments can be replaced or extended to enhance the potency of the ligands^22^.

**Fig. 1.**
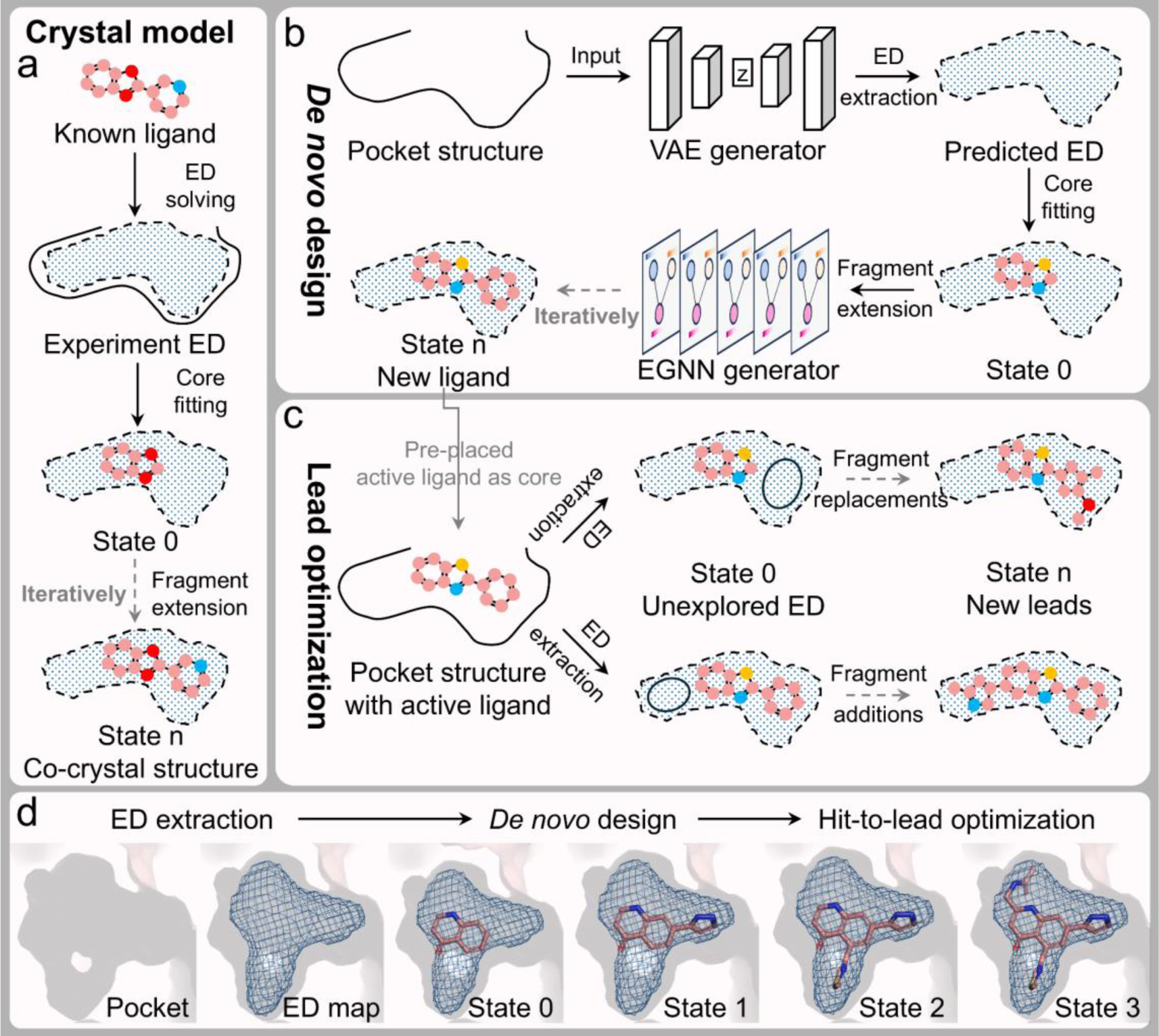
Architecture and workflow of ED2Mol. **a,** Illustration of the process by which structural biologists fit a ligand binding mode into an experimentally determined ED map, utilizing tools such as LigandFit. **b,** Schematic representation of the ED2Mol *de novo* design process. The ED for generated molecules is derived from the pocket structure via a VAE generator. A starting core is located and oriented near the largest contiguous region within the ED. Subsequently, fragments are iteratively extended using an EGNN generator, which is guided by the ED environment, enabling the molecule to evolve progressively from state 0 to state n. **c,** Schematic representation of the ED2Mol lead optimization process. A pre-placed active ligand serves as the initial core. Initially, unexplored electron density is identified at state 0, and the lead ligand is iteratively refined through the replacement or addition of fragments to occupy voids within the ED. **d,** An illustrative example showcasing the practical overall 3D molecular evolution.

In light of these observations, we propose ED2Mol, an ED-guided structure-aware DGM for *de novo* molecular design and lead optimization using the FBDD strategy. In Brief, ED2Mol leverages a variational autoencoder (VAE) to impute ligand ED from the pocket structure, portraying the space of “pseudo-ligands”. Subsequently, it applies a stratified peak density search algorithm to firstly fit the core, and iteratively extending fragments into unoccupied ED through an equivariant graph neural network (EGNN) (Fig. 1b). Thereupon, ED2Mol readily accommodates lead optimization by expanding a predefined core structure (active compound) into a complete lead molecule fragment-by-fragment, guided by the ED (Fig. 1c). These features make our method versatile, allowing for the generation of hit/lead compounds tailored to various application scenarios in drug design (Fig. 1d). More details of ED2Mol are described in the Methods and Supplementary Fig. 1.

Our comprehensive evaluation results demonstrate that ED2Mol outperforms the compared state-of-the-art (SOTA) DGMs across two pivotal tasks in drug discovery: *de novo* generation and lead optimization. For *de novo* generation task, regarding intermolecular reliability, ED2Mol remarkably surpasses other methods in generating reliable binding modes, making it highly credible for pharmacologists engaged in the design-make-test-analyze (DMTA) cycle. Regarding intramolecular reliability, ED2Mol also achieves the SOTA results in terms of topological plausibility, synthetic accessibility, and drug-likeness. In addition to benchmarking on the widely used Directory of Useful Decoys-Enhanced (DUD-E) dataset, which dominantly consists of 94 conserved orthosteric sites^23^, we also introduce a more challenging dataset, the AlloSteric Benchmark-Enhanced (ASB-E) dataset, comprising 112 diverse allosteric pockets. While other methods exhibit a noticeable performance decline on this dataset, ED2Mol maintains its robust superior performance. For lead optimization task, we present two representative case studies showing ED2Mol can be successfully integrated into lead design for inhibiting Ski2-like RNA helicase (Brr2) via fragment replacement and activating peroxisome proliferator-activated receptor gamma (PPARγ) via fragment growing strategies. Notably, ED2Mol were applied in four real-world drug design campaigns—two for *de novo* design and two for lead optimization. Wet-lab experiments validated the capability of ED2Mol to generate a diverse range of bioactive molecules, including both orthosteric and allosteric types, as well as inhibitors and activators, along with optimal binding modes for three important drug targets: fibroblast growth factor receptor 3 (FGFR3), cell division control protein 42 homolog (CDC42) and hexokinase-4 (GCK). Collectively, these extensive experiments underscore the effectiveness and practical utility of ED2Mol in hit/lead compound discovery.

## Results

### Model performance for *de novo* generation

A pocket-aware 3D DGM should effectively satisfy two design criteria crucial for drug discovery. First, from an intermolecular perspective, it should learn the optimal geometric distributions of ligands within binding pockets to generate binders with optimal binding modes. Second, from an intramolecular perspective, it should learn the pharmacologically relevant topological distributions of ligands across diverse pockets, as these distributions dictate essential pharmacological properties. To comprehensively evaluate ED2Mol against existing SOTA models regarding these two criteria, we not only employed the widely used DUD-E dataset but also introduced a more challenging, meticulously curated ASB-E dataset. For the first design criterion, we defined a reliable generated binder based on four aspects: binding site, efficiency, validity, and stability, as detailed below. For the second criterion, we assessed the pharmacological characteristics of the generated molecules, including synthesizability, drug-likeness, and structural diversity.

#### Benchmarks and baselines

The ED map offers a more intuitive and physically grounded understanding of ligand atomic structures compared to other expert-crafted rules, potentially enabling a more generalized depiction^16^. We hypothesize that incorporating such descriptors into DGMs enables them to learn more robust ligand-binding patterns and achieve stable generation results across diverse protein pockets. A comprehensive benchmark dataset that accounts for pocket diversity is essential for this assessment. However, we found that most DGMs are typically assessed using highly evolutionarily conserved endogenous pockets (also termed as orthosteric pockets)^23,24^. Rare attention has been paid to assessing their performance on non-canonical pockets, which predominantly include a broader range of less conserved allosteric pockets that are topologically distinct from orthosteric sites and can theoretically be located anywhere on the protein surface. Besides, allosteric pockets tend to be more superficial and smaller than orthosteric pockets, posing greater challenges for designing active compounds^25-27^.

To address these, two benchmark datasets were employed in this study. The first dataset, DUD-E, is widely used and comprises 94 pharmacological targets after retaining only orthosteric pockets. The second dataset, ASB-E, meticulously curates 112 high-quality and diverse allosteric pockets (see Methods for detail). The baselines in our study for comparison comprise five SOTA pocket-conditional DGMs, spanning from the atom-wise autoregressive models like Pocket2Mol^28^, GraphBP^29^, ResGen^30^, to fragment-wise autoregressive model FLAG^31^ to a non-autoregressive diffusion-based model TargetDiff^32^ (see Supplementary Methods for detail). For each pocket in both benchmark datasets, 1,000 molecules were generated using ED2Mol and each of the five baseline models, respectively.

#### Binding mode reliability of generated molecules

The primary objective of DGMs is to generate molecules that tightly bind to a given pocket in an optimal binding mode, which is interpreted through four perspectives: binding site, efficiency, validity, and stability. To begin with, despite the models are pocket-aware, the binding sites of the generated molecules may vary. We visualized the binding sites of 1,000 generated molecules within both orthosteric and allosteric sites of three representative drug targets: G-protein-coupled receptors (GPCRs), kinases, and nuclear receptors^33^. As illustrated in Supplementary Fig. 2, for proteins with diverse shapes, ED2Mol consistently generates molecules located within the inner regions of the respective pockets. Conversely, GraphBP produces molecules that are diffusely distributed around the pockets, with many located outside the pockets. Pocket2Mol and ResGen also primarily generate molecules within the target pocket, but their atomic density is more discrete and sparser relative to other models. This discretization likely arises from overlapping similar sub-structures, given that the two methods produce molecules with limited diversity (see data below). The atomic density of molecules generated by FLAG shows a dispersion from the pocket center to the surrounding area, possibly due to stepwise error accumulation and eventual structural collapse (Supplementary Fig. 3). TargetDiff performs well on some pockets but does not fully explore the space of allosteric pockets in the GPCR and nuclear receptor.

Next, DGMs have been observed to generate molecules with large sizes and bulky fused rings (Supplementary Fig. 3), which can potentially inflate computational binding affinity estimates^34^. To alleviate this affinity bias and allow smaller compounds with moderate affinity to be also attractive for subsequent lead optimization, we employed ligand efficiency (LE), which scales binding affinity by molecular size^35,36^. The generation success rate, commonly defined as the percentage of generated molecules with a LE superior to that of the reference co-crystal ligands, is utilized for evaluation. As demonstrated in Fig. 2a and Table 1, ED2Mol outperforms all other SOTA models in terms of LE, achieving success rates of 67.3% on the DUD-E dataset and 68.9% on the ASB-E dataset, surpassing other models by an average of 6.5% to 61.3%. This superior performance indicates that ED2Mol generates molecules with efficient binding modes across different pocket types. It is abnormal that ResGen and GraphBP produce poor binding molecules that were scored higher-than-zero. A careful examination of the generated molecules revealed that a substantial proportion suffer from irrational binding sub-structures and steric clashes with receptor atoms (see data of binding validity). Besides, as mentioned above, molecules generated by GraphBP are distributed randomly, with some even located outside the protein surface, leading to even weaker binding interactions. It is worth noting that TargetDiff, Pocket2Mol and ResGen exhibited a notable decrease in LE on the ASB-E dataset compared to the DUD-E dataset, whereas ED2Mol maintained robust performance across both datasets. This robustness to diverse, unseen allosteric pockets likely stems from its fragment-based protocol, which generates fragments that bind more efficiently to small, superficial allosteric pockets, a characteristic also contributing to FLAG’s consistent performance. On the other hand, ED2Mol’s ability to capture external complementary shape and internal precise electron distribution of ED map, further enhances its ligand-binding efficiency.

**Fig. 2.**
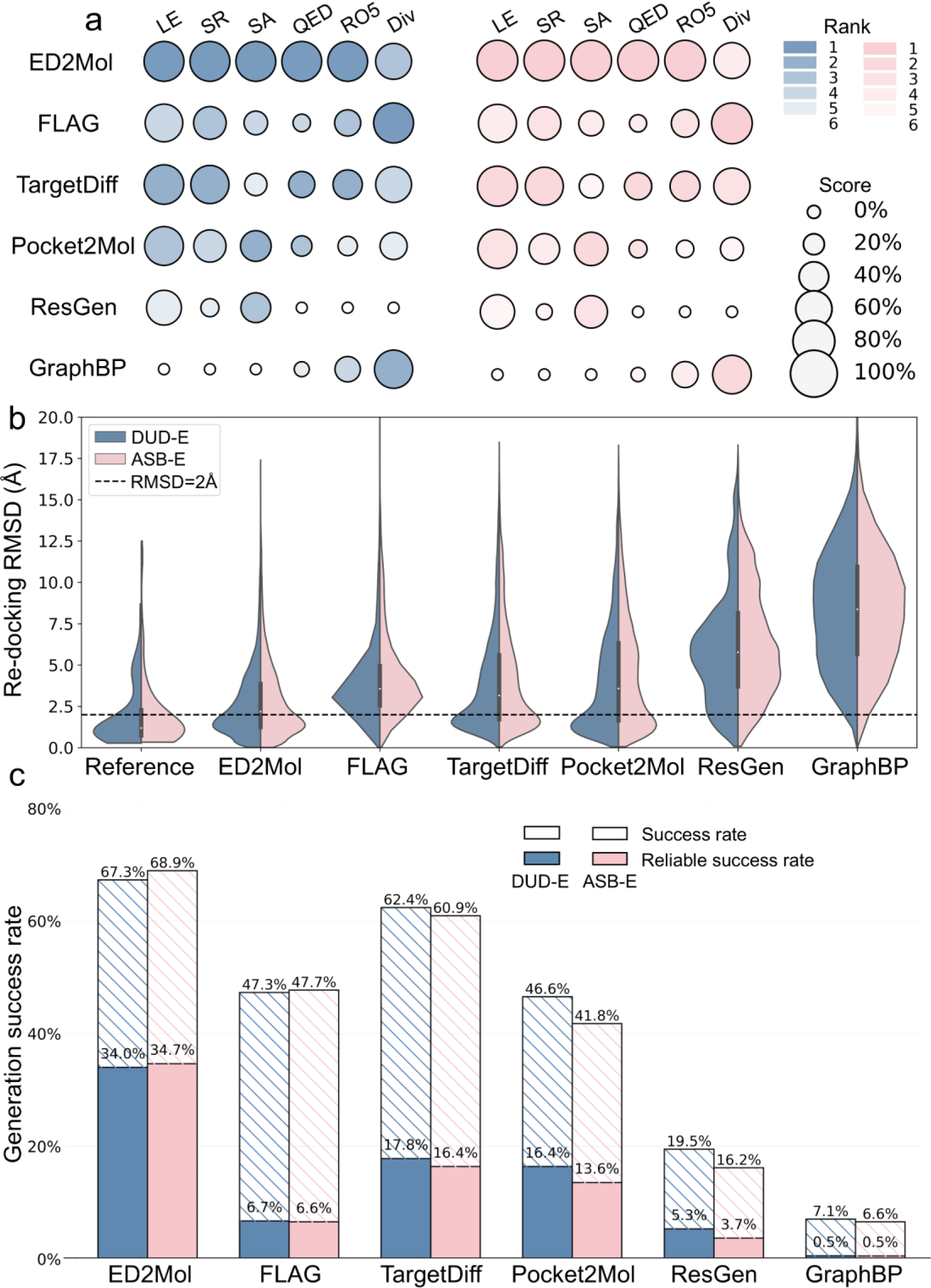
Performance comparison of DGMs using the DUD-E (blue) and more challenging ASB-E (red) sets. **a,** Metrics encompass ligand efficiency (LE), success rate (SR), synthetic accessibility (SA), quantitative estimation of drug-likeness (QED), Lipinski’s Rule of Five (RO5), and molecular diversity (Div). The size of circles corresponds to the mean normalized score for the respective metric (from 0% to 100%), while the darkness of circles reflects the performance ranking (from 1 to 6). The original data are provided in Table 1 for LE and SR, and in Supplementary Table 3 for the remaining. **b,** Distribution of RMSD between the original generated poses and re-docked poses. The reference violin plot is derived from re-docking the ligand from the co-crystal structure. **c,** Comparative analysis of generation success rates between ED2Mol and baselines. The striped bars (success rate), indicating the proportion of generations with LE higher than the reference ligand, while the solid bars (reliable success rate) signify the proportion of generations that also meet the binding validity and stability criteria.

**Table 1.** Binding metrics from multiple perspectives on DUD-E and ASB-E sets.

| Datasets | Methods | Binding efficiency |  | Binding validity | Binding stability |
| --- | --- | --- | --- | --- | --- |
| | | LE (↓) | SR (↑) % | PB-validity (↑) | RMSD (↑) % < 2Å |
| DUD-E<br>(94 targets) | ED2Mol | **-0.36** | **67.3** | **97.5** | **47.5** |
|  | FLAG | -0.22 | 47.3 | 58.4 | 16.2 |
|  | TargetDiff | -0.33 | 62.4 | 58.4 | 33.4 |
|  | Pocket2Mol | -0.29 | 46.6 | 62.8 | 34.8 |
|  | ResGen | -0.08 | 19.5 | 31.3 | 11.2 |
|  | GraphBP | 0.83 | 7.1 | 31.3 | 1.8 |
| ASB-E<br>(112 targets) | ED2Mol | **-0.35** | **68.9** | **97.0** | **44.9** |
|  | FLAG | -0.20 | 47.7 | 60.3 | 13.3 |
|  | TargetDiff | -0.31 | 60.9 | 57.7 | 29.0 |
|  | Pocket2Mol | -0.26 | 41.8 | 63.3 | 28.0 |
|  | ResGen | -0.02 | 16.2 | 36.8 | 7.6 |
|  | GraphBP | 0.79 | 6.6 | 33.1 | 1.6 |

Furthermore, it was discovered that many generated molecules are invalid, we hence conducted a comprehensive binding validity examination utilized 18 criteria outlined in PoseBusters^12^. These criteria evaluate the intramolecular and intermolecular validity of the generated binding poses. Ligand poses that pass these criteria are termed PB-valid. ED2Mol substantially outperforms other models in generating high-quality, PB-valid binding poses, as shown by the following average rankings on both datasets (Supplementary Table 1 and 2): ED2Mol (97.3±0.4%) > Pocket2Mol (63.1±0.4%) > FLAG (59.4±1.4%) > TargetDiff (58.1±0.5%) > ResGen (34.1±3.9%) > GraphBP (32.2±1.2%). In specific, ED2Mol demonstrated the highest feasibility, with only one-fortieth of its generated poses encountering protein-ligand clashes on average. In contrast, in terms of intermolecular validity, over half of the molecules generated by ResGen and GraphBP suffer from steric clashes, while this ratio is nearly one-fifth for Pocket2Mol and FLAG, and one-tenth for TargetDiff. In terms of intramolecular validity, all baseline models encountered distorted geometries. FLAG and TargetDiff primarily struggle with predicting correct bond angles, whereas Pocket2Mol and ResGen often fail to predict valid bond lengths, with GraphBP performing poorly in both respects.

An additional observation is that many of the generated ligand binding poses undergo dramatic rearrangements after re-docking, suggesting that DGMs may not capture the globally optimal interactions, thereby impeding the selection of drug candidates by pharmacologists^11,37^. To investigate binding stability, we conducted re-docking experiments comparing the original poses to re-docked poses, calculating the minimum root-mean-square-deviation (RMSD) across the top 10 docking poses. Remarkably, reference co-crystal ligands successfully recovered 72% of their crystallized poses (RMSD < 2Å) across both datasets, validating the use of re-docked poses, at least in both benchmark datasets, as a surrogate ground truth for evaluating the fitness of directly generated poses. When we re-docked molecules generated by ED2Mol and the baselines following the same pipeline, ED2Mol showed the highest fidelity in maintaining binding poses (Fig. 2b and Table 1), with 12.7% more stable poses than the second-best model, Pocket2Mol, on the DUD-E dataset (34.8% total) and 15.9% better stability than the second-best model, TargetDiff, on the ASB-E dataset (29.0% total).

Collectively, a reliable binding mode for generated ligands should be characterized by greater binding efficiency (success rate), high-quality (PB-valid) and stable (RMSD < 2Å upon re-docking) binding pose. Based on these three criteria, we define the reliable success rate as the proportion of generated ligands with reliable binding modes (Figure 2c). ED2Mol achieved a reliable success rate of 34.4±0.5%, more than twice as high as the best-performing baseline models, TargetDiff (17.1±1.0%) and Pocket2Mol (15.0±2.0%), averaged across both datasets. FLAG (6.7±0.1%) and ResGen (4.5±1.1%) produced very few reliable ligands, and GraphBP produced almost no reliable ligands in our evaluation.

#### Pharmacologically desirable properties of generated molecules

Beyond reliable binding modes, pharmacologists prioritize ligands with favorable pharmacological properties, such as ease of synthesis and drug-likeness when selecting compounds for further DMTA cycles^38,39^. For this purpose, we accessed the generated molecules based on three widely recognized metrics: synthetic accessibility (SA), drug-likeness (QED), and adherence to Lipinski’s Rule of Five (RO5). ED2Mol obtains the best scores across all three metrics (Supplementary Table 3 and Fig. 2a), highlighting its capability to generate pharmacologically desirable molecules that are both readily synthesizable and drug-like. Additionally, molecules produced by ED2Mol show comparable diversity to those produced by the baselines.

### From *de novo* design to lead optimization

Currently, it is virtually impossible to directly generate fully optimized potential drugs through automated de novo design using DGMs^40^. One of the primary reasons for this limitation is the reliance on experimentally determined crystallographic data or computationally docked data (e.g., CrossDocked2020 dataset^24^) for training, which do not account for protein flexibility. Such limitations are understandable, as incorporating pocket flexibility would significantly increase computational demands and generation time. Indeed, the binding pockets derived from crystallographic data represent only a single, static druggable conformation, while in reality, they are highly dynamic and exist in multiple druggable conformations^41^. Furthermore, ligand loading can influence these dynamics via conformational selection and induced-fit mechanisms^42,43^. Nevertheless, most DGMs assume that the ligand generation process does not alter the binding pocket geometry. As a result, once the complex structure of the protein and ligand is experimentally determined, opportunities may arise to optimize the currently generated compounds to achieve higher potency (i.e., lead compounds). Our subsequent wet-lab validation of the generated molecules against cancer target FGFR3 corroborated this observation.

Therefore, lead optimization constitutes a crucial step toward fully *de novo* drug design. A primary strategy involves elaborating known active ligands by swapping and/or adding new moieties (fragments) to establish additional interactions with the receptor. This strategy is directly applicable to ED2Mol through a fine-tuned workflow, which bypasses the core searching procedure and instead utilizes a pre-located core scaffold as a template for further fragment elaboration. To demonstrate the utility of ED2Mol in identifying effective fragment replacements or additions, we present two representative practical case studies from the literature: fragment replacements of Brr2 inhibitors and fragment additions of PPARγ activators. The baselines employed in this experiment range from autoregressive fragment-wise FLAG to the diffusion-based DiffDec^44^ and the traditional method, FragGrow^45^. For a fair comparison, FLAG was adjusted following the approach of Xie et al. for lead optimization tasks^44^ (see Supplementary Methods for detail). For each task, we generated 3,000 compounds using ED2Mol and the baseline methods, respectively.

#### Design of Brr2 inhibitors via fragment replacements

As a central component of the spliceosome, the Ski2-like RNA helicase Brr2 plays a critical role in the precise activation, catalysis, and disassembly of the spliceosome and ensure the fidelity of gene expression, rendering Brr2 an attractive genetic drug target^46^. Iwatani-Yoshihara et al. reported a potent inhibitor targeting the allosteric pocket of Brr2^47^. Initially, they identified a hit compound (compound **3**) that bound to the allosteric pocket of Brr2 through high-throughput screening, followed by co-crystal structure determination (PDB ID: 5URJ). This structure revealed that the core scaffold of compound 3, 6-benzyl-4,6-dihydropyrido[4,3-d]pyrimidine-2,7(1H,3H)-dione, forms a key hydrogen bond interaction with residue I1681. By retaining the core scaffold and substituting the remaining fragment, they developed lead compound **9**, which exhibited a 67-fold increase in inhibitory activity.

To better emulate practical scenarios where prior knowledge of the lead co-crystal structure is unavailable, we utilized the receptor pocket induced by hit compound **3** (PDB ID: 5URJ). Similarly, the core scaffold 6-benzyl-4,6-dihydropyrido[4,3-d]pyrimidine-2,7(1H,3H)-dione was scissored out as the starting point and ED2Mol was integrated for fragment replacements. The ligand ED predicted by ED2Mol successfully explored vacant regions within the sub-pocket encompassing residues T1197, F1255, P1257, K1716, and F1713. Remarkably, among all methods tested, only ED2Mol was able to recover lead compound **9**. Meanwhile, the binding pose generated by ED2Mol closely aligned with that predicted by molecular docking (RMSD = 0.56 Å), underscoring its capability to produce reliable binding modes (Fig. 3b).

**Fig. 3.**
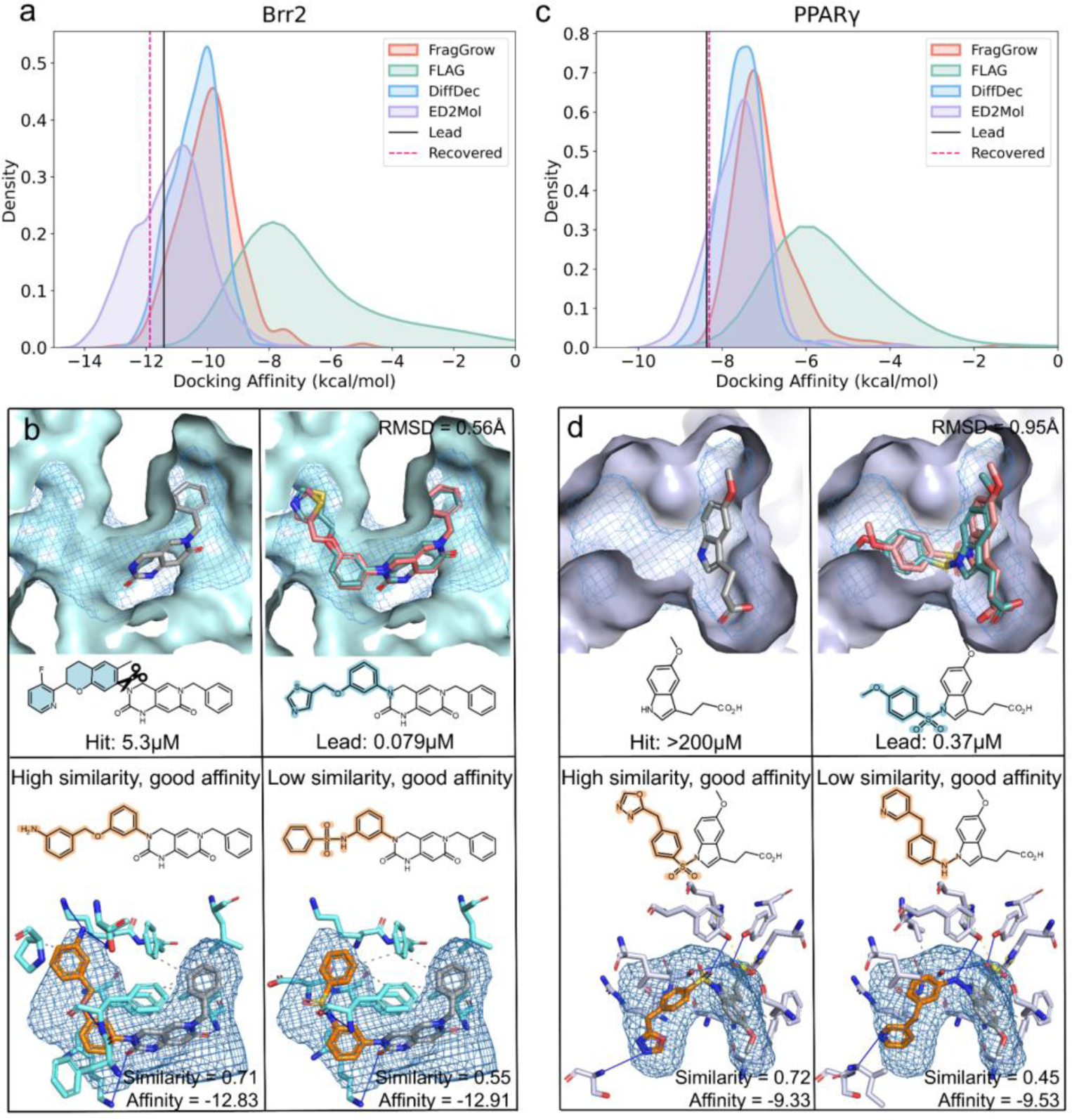
Retrospective Lead optimization case studies. Left, fragment replacements of Brr2 inhibitors. Right, fragment additions of PPARγ activators. **a** and **c,** Distribution of docking affinity scores for optimized samples of Brr2 (**a**) and PPARγ (**c**) generated by ED2Mol and other methods. The black solid lines denote the scores of computationally re-docked (for Brr2) and experimentally determined (for PPARγ) lead compounds. The red dashed lines represent the scores of compounds generated by ED2Mol that recover the lead compounds. **b** and **d,** In the upper panels, the reported lead compound (upper right, in red) can only be recovered by ED2Mol (upper right, in blue) from the original hit compound (upper left, in gray). The elaboration moieties involved are highlighted in blue. In the bottom panels, two representative samples with superior affinity compared to the reported lead compound are shown: one with high chemical similarity to the reported lead, and the other with low similarity. The diverse elaboration moieties are highlighted in orange. Intermolecular interactions are analyzed using PLIP.

In addition, we assessed the generated molecules using computational binding affinity scores, as affinity, rather than LE, is critical during the lead optimization stage. The score distribution suggests that molecules generated by ED2Mol not only score better than those generated by baselines but also that some achieve superior scores to the lead compound **9** (Fig. 3a). A closer inspection revealed two distinct groups of molecules (Fig. 3b). The first group, termed the “high similarity & good affinity” group, closely resembles compound **9** in molecular topology (Fingerprint similarity = 0.71) but binds more tightly to the pocket. This improvement was attributed to additional interactions: it preserves the hydrogen bond between the oxygen atom of the swapping fragment and the main chain amine of residue F1255, as seen in compound **9**, while its terminal amino nitrogen forms additional hydrogen bonds with residues T1197 and K1716 in the sub-pocket. The second group, referred to as the “low similarity & good affinity” group, presents novel scaffolds. For example, the nitrogen of a sulfonamide group in the representative molecule maintains polar interactions with F1255, while the phenyl ring establishes more extensive non-polar interactions with surrounding residues. These findings demonstrate that ED2Mol can effectively identify moiety replacements that optimize diverse and favorable interactions within the protein pocket.

#### Design of PPARγ activator via fragment additions

PPARγ activators represent a promising therapeutic strategy for treating type 2 diabetes and associated metabolic syndromes^48^. Through biochemical screening, Artis et al. identified potent PPARγ activators, compound **1**, and solved its co-crystal structure (PDB ID: 3ET0)^49^. Utilizing a structure-guided fragment growing strategy, they obtained lead compound **3** (PDB ID: 3ET3), which exhibited over a 70-fold increase in potency for PPARγ.

Herein, we investigate the ability of ED2Mol to detect sub-pockets and optimize a more challenging activator compound. Using the co-crystal structure of compound **1** as input, ED2Mol captured additional growing space surrounding residues R288, Y327, L330, and S342. ED2Mol was the only method to recover lead compound **3** with a reliable binding pose, consistent with the lead-bound complex structure (RMSD = 0.95 Å) (Fig. 3d). Following our previous experiments, we computationally scored the binding affinity of the generated molecules. The binding affinity distribution of ED2Mol-generated molecules was more left-skewed compared to both experimentally confirmed lead compounds and other methods (Fig. 3c). Likewise, the representative ligand in the “high similarity & good affinity” group retains the hydrogen bond with the hydroxyl group of residue Y327 via the same sulfonyl group as compound **1**. Instead, the ligand in the “low similarity & good affinity” group forms this interaction via an alternative amide group. Moreover, both ligands gained additional polar interactions with the main chain of residue S342 and non-polar interactions with residues R288, L330 and others, accounting for the enhanced binding potency observed in both groups (Fig. 3d).

### From benchmarking evaluation to wet-lab verification

To substantiate the effectiveness of ED2Mol, we conducted wet-lab experiments on four real-world scenarios: FGFR3 orthosteric inhibitors generation and optimization, CDC42 allosteric inhibitors *de novo* generation, and GCK allosteric activators lead optimization. Despite these proteins are well-established targets for various diseases, particularly in anti-tumor and anti-diabetic therapies, there remains a paucity of efficient small-molecule intervention agents. Expanding the chemical space to identify potent compounds thus holds great promise.

#### FGFR3 orthosteric inhibitors generation and optimization

FGFR3, a member of the tyrosine kinase superfamily, propagates FGF-related signals that regulate numerous biological processes, including cell proliferation, differentiation, and survival^50^. Dysregulation of FGFR3 is implicated in a wide range of diseases, notably cervical and bladder carcinomas^51,52^. Blocking the dysregulated FGFR3 signaling has been established as a promising therapeutic approach for these malignancies.

To address this, ED2Mol was employed to generate novel FGFR3 inhibitors. The co-crystal structure of FGFR3 bound to an ATP analog (PDB ID: 4K33) was used to define the orthosteric binding pocket as the target site (Fig. 4a)^53^. From scratch, ED2Mol generated 50,000 molecules within 100 minutes on a laptop equipped with a GeForce RTX 4080 GPU. It was shown that effective orthosteric kinase inhibitors are expected to exhibit both favorable pharmacological properties and a binding mode that mimics the interaction pattern of ATP’s nucleobase, forming hydrogen bonds with the kinase hinge region^54,55^. Hence, to narrow the initial pool of candidates, the following criteria were applied: (1) molecular weight > 250, (2) compliance with RO5, (3) QED > 0.8, (4) SA score < 4.5 and (5) at least one hydrogen bond with the hinge region. These filters reduced the pool to 687 molecules. Typically, to minimize redundancy (i.e., highly similar molecules), the molecules were clustered based on 2D structural similarity, and 5 representative candidates were then selected (Supplementary Fig. 4). Among these, **F4** exhibited the closest hinge-binding pattern to ATP, establishing hydrogen bonds with the backbones of residues E556 and A558, while its fluorobenzene moiety engaged in hydrophobic interactions with L478, V486, N562, and L624. **F4** was thus prepared for wet-lab validation, with synthetic routes (Scheme 1) provided in the Supplementary Information. The surface plasmon resonance (SPR) assay confirmed that **F4** reversibly interacted with FGFR3 in a dose-dependent manner, yielding an equilibrium dissociation constant (K_D_) of 599.0 μM (Fig. 4b and Supplementary Fig. 5).

**Fig. 4.**
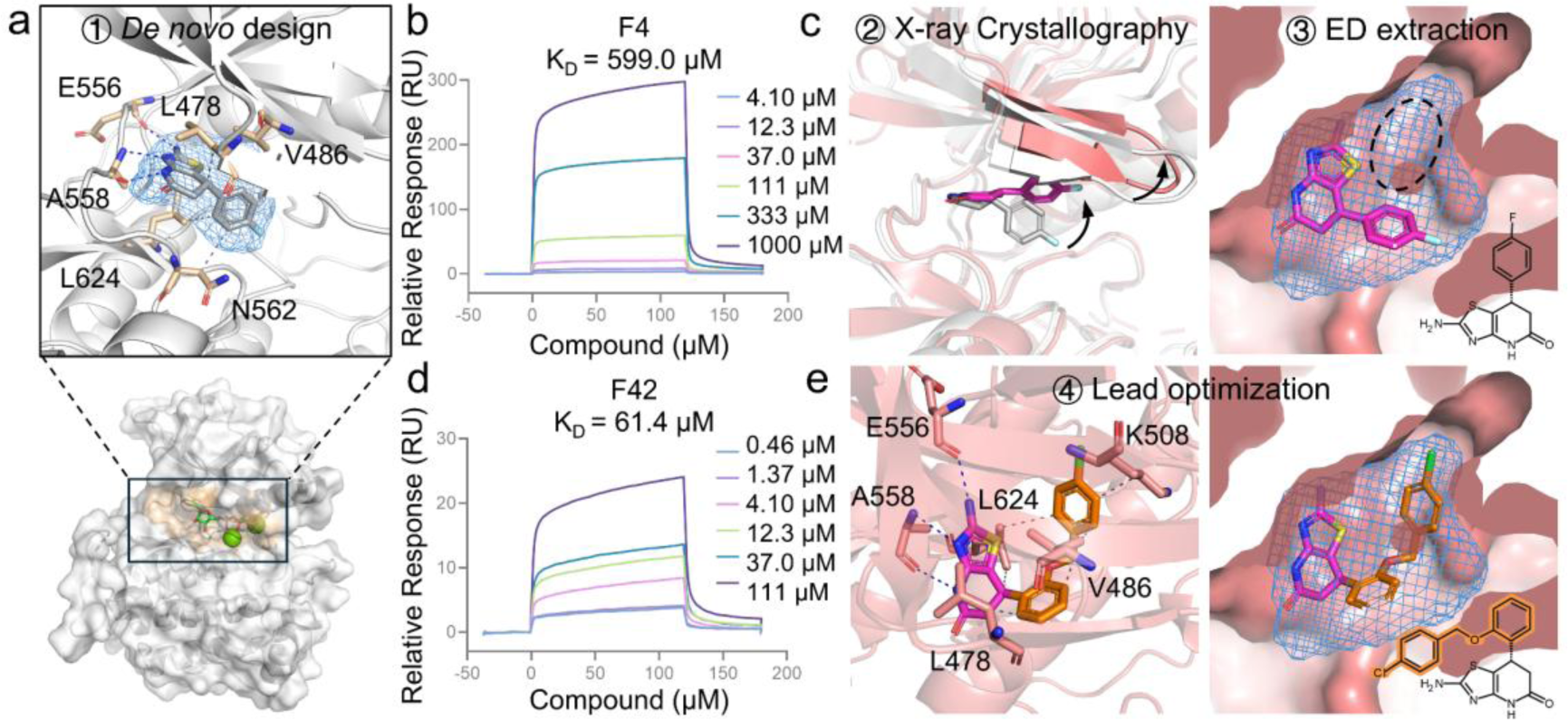
Discovery of new FGFR3 orthosteric inhibitors through practical applications of ED2Mol. **a,** The generated F4 compound at the orthosteric pocket of FGFR3. Bottom, the input structure (PDB ID: 4K33) of FGFR3 (gray surface) in complex with orthosteric ligand (green stick). Upper, a magnified view of the generated binding mode of F4 compound, where the 2-amino-6,7-dihydrothiazolo[4,5-cb]pyridin-5(4H)-one moiety forms critical hydrogen bonds with hinge residues E556 and A558. **b,** Dose-dependent curve for F4 measured by SPR assay. **c,** Co-crystal structure of FGFR3 in complex with F4 (PDB ID: 9KFU). Left, superimposition of the binding mode of F4 with FGFR3 as generated by ED2Mol (gray) and that determined experimentally by X-ray crystallography (magenta). Arrows indicate conformational changes in the Gly-rich loop and adjacent β1- and β2-strands, as well as the fluorobenzene moiety of F4 compound. Right, based on the newly resolved complex structure, the second-round ED extraction captures an additional sub-pocket, indicated by the black dashed circle. **d,** Dose-dependent curve for F42 measured by SPR assay. **e,** Lead optimization leading to the F42 compound. Left, FGFR3-F42 interaction analysis as determined by PLIP. Right, the binding mode of the generated F42 fully occupies the newly identified sub-pocket, with the chemical structure shown at the bottom, and the swapped portion highlighted in orange.

To elucidate the true molecular interactions between the **F4** and FGFR3, the co-crystal was resolved via X-ray crystallography at a high resolution of 1.4 Å. The experimentally determined binding mode closely matched that direct generated via ED2Mol (RMSD = 2.00 Å). Intriguingly, the fluorobenzene moiety in the co-crystal structure adopted an upward orientation, driving conformational changes in the Gly-rich loop as well as adjacent β1- and β2-strands, leading to a widened orthosteric site. This observation is consistent with the induced-fit binding mechanism frequently observed in kinase inhibitors, offering new opportunities for the further optimization of F4^56^ (Fig. 4c).

Indeed, capitalizing on the **F4**-induced conformational changes in FGFR3, ED2Mol captured an extended sub-pocket near residues L624 and K508. Subsequently, following the protocol previously used for Brr2 inhibitors, lead optimization was performed through fragment replacements while retaining the essential hinge-binding moiety of **F4**. We produced 10,000 molecules by ED2Mol on a single GeForce RTX 4080 GPU in 20 minutes. Filtering criteria were adjusted to: (1) molecular weight > 250, (2) compliance with RO5, (3) QED > 0.7, (4) SA score < 4. After these filters, 361 molecules remained and were clustered into 3 groups (Supplementary Fig. 6). From these, candidate **F42** was chosen based on its relatively facile synthetic route (Scheme 2 in Supplementary Information) recommended by SA score (3.6) and medical chemists. The SPR assay determined that **F42** possessed a KD of 61.4 μM, representing a 9.8-fold increase in potency compared to the initial **F4** compound (Fig. 4d and Supplementary Fig. 7). This improvement can be largely attributed to F42’s entire occupation of the newly resolved sub-pocket, promoting additional contacts with the surrounding residues (Fig. 4e).

### CDC42 allosteric inhibitors *de novo* generation

CDC42, a small GTPase, functions as a binary switch in a variety of downstream signaling pathways, including cell migration and angiogenesis, via cycling between its active GTP-bound state and inactive GDP-bound state^57,58^. Overexpression of CDC42 is frequently observed in multiple human cancers, where it promotes tumor progression, rendering it an attractive target for anti-tumor therapies^59^. Small-molecule inhibitors of CDC42 have the potential to serve as therapeutic agents against these malignancies. Recently, Jahid and co-workers identified a compelling allosteric pocket located at the protein-protein interaction (PPI) site between CDC42 and its downstream partner p21-activated kinase (PAK), which poses a new avenue for the design of allosteric inhibitors^60^.

Again, to discover new active inhibitors, we deployed ED2Mol to generate molecules targeting the allosteric PPI site taking the CDC42-PAK (PDB ID: 2ODB) crystal structure as input (Fig. 5a). First, ED2Mol generated a set of 50,000 molecules. Second, these molecules were filtered using four criteria: (1) molecular weight > 250, (2) compliance with RO5, (3) QED > 0.8, (4) SA score < 4.5. This filtering process eliminated molecules with low drug-likeness and poor synthetic feasibility, resulting in 1,547 candidates that satisfied all conditions. Third, these candidates were clustered into 5 distinct groups, from which 5 representative molecules were selected (Supplementary Fig. 8). Among the representatives, compound **C1** exhibited a stronger π-π stacking interaction with residue F37 and was deemed more synthetically accessible. Again, the superimposition of the binding mode produced by ED2Mol and that derived by molecular docking reveals a considerable similarity in the binding mode (RMSD = 0.82 Å), reinforcing the notion that ED2Mol can generate reliable predictions. Following the synthetic routes detailed in the Supplementary Information (Scheme 3), **C1** was synthesized and subjected to bioactivity testing. It demonstrated inhibitory activity against CDC42, with a half-maximal inhibitory concentration (IC_50_) value of 47.58 ± 3.71 μM (Fig. 5b). In addition, we synthesized **C11** (Scheme 4), a close structural analogue of **C1** within the same cluster, to preliminarily explore the structure-activity relationship (SAR) of the symmetrical chlorobenzene of **C1**. The compound **C11** exhibited a reduced potency, with an IC_50_ value of 111.63 ± 0.90 μM, suggesting a comprehensive optimization is required to improve the potency of **C1** (Supplementary Fig. 9). Nevertheless, **C1** could serve as a valuable starting hit compound for subsequent investigations.

**Fig. 5.**
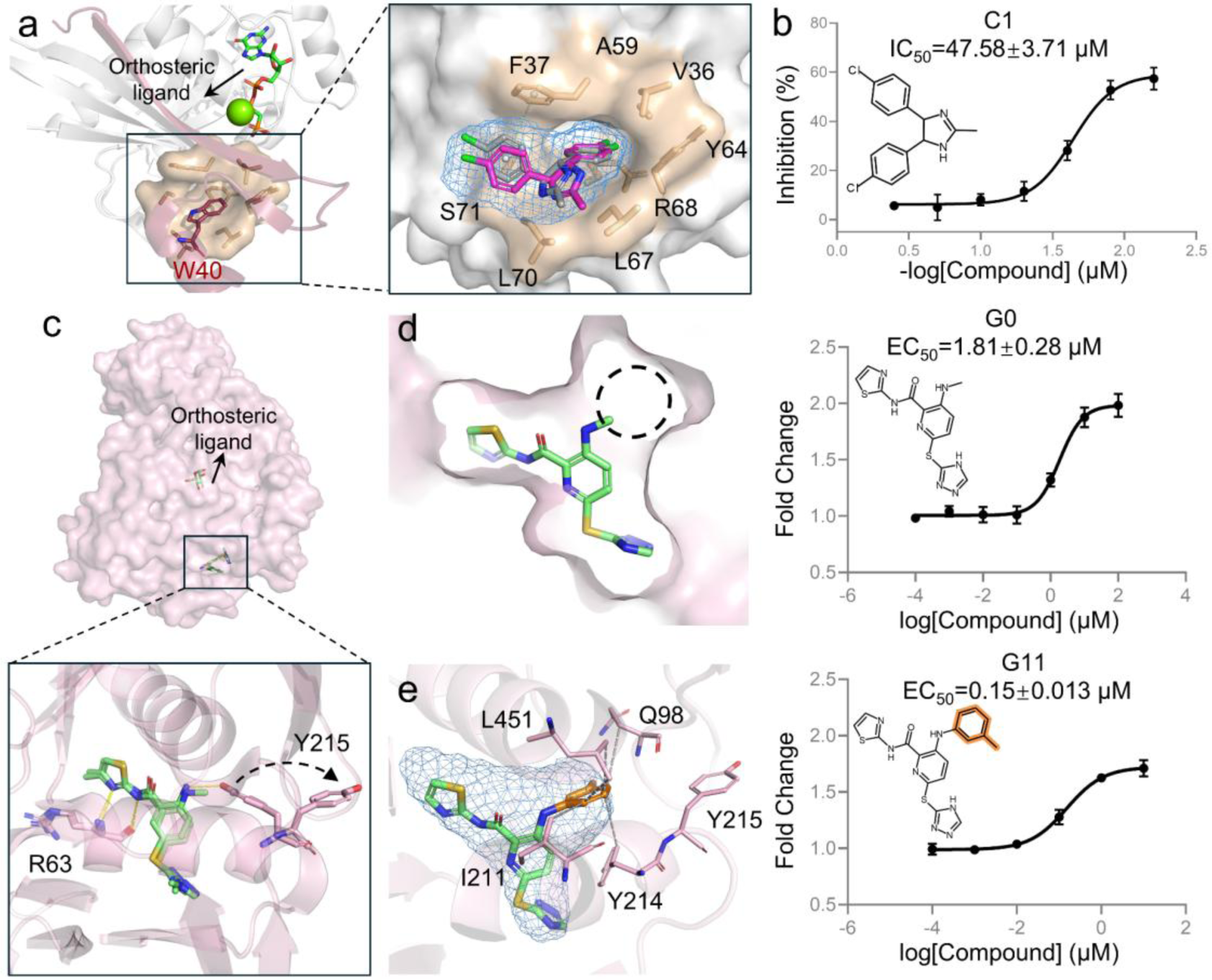
Discovery of new CDC42 allosteric inhibitors and GCK allosteric activators through practical applications of ED2Mol. a, The generated C1 compound at the allosteric pocket of CDC42. Left, the input structure (PDB ID: 2ODB) of CDC42 (gray cartoon) in complex with orthosteric ligand and allosteric PAK6 (red cartoon). The W40 (red)-mediated hydrophobic allosteric PPIs with surrounding residues (tint) are highlighted. Right, superimposition of the binding mode of C1 with CDC42 as generated by ED2Mol (gray) and that predicted by molecular docking (magenta). b, Dose-dependent curve for C1 measured by BRET assay. The experiments were performed independently three times. All data are shown as mean ± s.d. c, Two previously identified activators at the allosteric pocket of GCK. Upper, the input structure (PDB ID: 3GOI) of GCK (pink surface) in complex with orthosteric ligand and allosteric activator. Bottom, a magnified view of the superimposed binding modes of the activators, which shows that the added methyl group on the aniline NH group induces rotamer changes in Y215, as indicated by the black dashed arrow. d, Binding mode of hit compound G0. Left, ED extraction reveals an additional sub-pocket, as indicated by the black dashed circle. Right, Dose-dependent curve for G0 measured by DEC assay. The experiments were performed independently three times. All data are shown as mean ± s.d. e, Binding mode of the generated lead compound G11. Left, GCK-G11 interaction analysis suggests the grown methylbenzene group forms several hydrophobic interactions with residues Q98, I211, Y214 and L451. Right, Dose-dependent curve for G11 measured by DEC assay. The experiments were performed independently three times. All data are shown as mean ± s.d.

#### GCK allosteric activators lead optimization

GCK is an enzyme that catalyzes the phosphorylation of glucose to glucose-6-phosphate, playing an important role in glucose metabolism and glucose-dependent insulin secretion^61^. Small-molecule activators of GCK have garnered attention as therapeutic agents for type 2 diabetes due to their ability to suppress glucose production and stimulate insulin secretion^62^. For instance, using high-throughput screening campaign followed by X-ray crystallography, Nishimura, T. et al. led to the discovery of a sub-micromolar activator with a thiazole-amide scaffold, which activates GCK by binding to an allosteric pocket (PDB ID: 3FR0)^63^. Interestingly, further structural analysis revealed that a methyl addition on the aniline moiety could push the side chain of residue Y215 reoriented outwards the pocket, enabling the emergence of a new sub-pocket (PDB ID: 3GOI)^64^ (Fig. 5c).

Building on this observation, fragment additions were anticipated to enhance potency by complementarily occupying the newly formed sub-pocket. Similar to the fragment additions strategy for PPARγ activators, we used the co-crystal structure (PDB ID: 3GOI) as the original template. The central benzene was varied with pyridine, based on the suggestion by Mitsuya et al. that this modification could stabilize the methyl orientation through an intramolecular hydrogen bond with the nitrogen atom of the pyridine ring and the amide group, to build the binding mode of analog **G0**^64^. ED2Mol was set to initially propose 10,000 molecules based on the **G0** compound (Fig. 5d). To narrow down the selection, four filters were implemented: (1) molecular weight > 250, (2) compliance with RO5, (3) QED > 0.4, (4) SA score < 4. These filters yield 995 molecules, leaving 3 categories after clustering (Supplementary Fig. 10). From these, compound **G1** was preferred due to its ease of synthesis and effective occupation of the sub-pocket. The synthetic routes and molecular characterization of **G1** are provided in the Supplementary Information (Scheme 6). Bioassays demonstrated that **G1** achieved an improved potency with an EC_50_ of 290 nM, representing a 6.1-fold enhancement over **G0** (Supplementary Fig. 11). Again, a preliminary survey of SAR was conducted by preparing **G11** (Scheme 7), belonging to the same category as **G1**. **G11** showed even greater potency, with an EC_50_ of 150 nM, representing an 11.9-fold improvement over the original hit compound **G0** (Fig. 5e). While further SAR investigations are warranted, these bioassay results suggest that **G11** could be a strong candidate for later evaluation and development.

## Discussion

In this study, we describe ED2Mol, leveraging ED map as generative cues for highly effective and reliable structure-based *de novo* design and lead optimization. ED2Mol encompasses several desirable features that facilitate the practical development of potential drug candidates through FBDD strategies.

We evaluated ED2Mol across multiple benchmarking scenarios. In these evaluations, ED2Mol achieved consistently superior performance to other DGMs in terms of generation success rates, plausible intramolecular topologies and credible intermolecular interactions. It also exhibited adaptability to more challenging, unseen allosteric pockets. It is worth noting that over 75% of generated molecules by other DGMs exhibit improper or sub-optimal binding modes, severely compromising the reliability of the generated samples, regardless of their promising computational docking or drug-likeness scores. On the contrary, ED2Mol consistently produces reliable molecules, showing more than a 2-fold improvement in this regard. Besides, ED2Mol habitually produces molecules with better synthetic accessibility and drug-likeness profiles, thereby instilling greater confidence in subsequent stages of wet-lab chemical synthesis and bioactivity verification.

Furthermore, ED2Mol’s capabilities extend beyond *de novo* design to lead optimization. In the retrospective experiments, ED2Mol not only captures the unexplored regions within the pocket but also stands as the only model to successfully recover experimentally validated lead compounds, whether designing inhibitors or more challenging activators. Additionally, ED2Mol could identify alternative moieties and that may bind more effectively to the target site.

More importantly, the practical utility of ED2Mol has been validated through various wet-lab experiments. ED2Mol was utilized to design *de novo* or optimize small-molecule intervention agents for FGFR3, CDC42, and GCK, which are appealing drug targets for a range of diseases, particularly cancer and diabetes. These real-world scenarios encompass diverse binding pocket types, including both orthosteric and allosteric sites, and involve different ligand types, such as inhibitors and activators, demonstrating the robustness and generalizability of ED2Mol across diverse drug discovery contexts. As a result, ED2Mol identified two FGFR3 orthosteric inhibitors, two CDC42 allosteric inhibitors, and two GCK allosteric activators, all of which are valuable for further systematic research. Of note, the generated binding modes of these active compounds closely resemble those predicted by computational docking, and experimental co-crystallography further confirms the true molecular interactions between the active compound and the protein pocket.

Collectively, ED2Mol exhibits excellent performance across all the designed experiments, providing a versatile and reliable tool for the discovery and optimization of novel therapeutic agents. However, several avenues remain open for further enhancement. As previously mentioned, improvements could be achieved by incorporating protein conformational flexibility, exploring cryptic adjacent sub-pockets, and generating molecules based on conformational ensembles^65^. Moreover, one may optimize binding affinity or other pre-clinical desirable pharmalogical properties (e.g., low toxicity) individually or concurrently through explicit guidance or reinforcement learning^66,67^.

## Methods

### Details of ED2Mol

Inspired by LigandFit, ED2Mol adopts a similar fragment-wise sampling strategy comprising three main components: (1) extracting the ligand ED from pocket structure; (2) placing cores within the ED and (3) generate a molecule by iterative fragments extension into the ED.

#### Ligand ED extraction

Constructing an ED map for a crystal structure involves first calculating the structure factors for the reflections in reciprocal space and then applying an inverse Fourier transform to convert these factors into a real-space map. According to Bragg’s Law, the diffraction pattern of a crystal reflection can be characterized by a series of parallel, hypothetical crystalline planes that mathematically intersect the axes of the crystal unit cell. These planes are described by Miller indices (*hkl*), which correspond to the reciprocals of the intercepts along the unit cell axes. Thereafter, the structure factor, *F*(*hkl*), can be obtained by summing the contributions of waves scattered by *N* atoms in the crystal along the direction of *hkl* reflection:

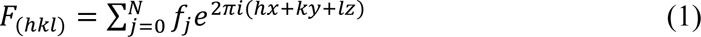

where each term involves the atomic scattering factor *f*_*j*_, representing the X-ray scattering power of the atom *j*. Once the structure factors are known, the ED *ρ*_*xyz*_ at any point (*x*, *y*, *z*) in the unit cell can be computed via the inverse Fourier transform *F*(*hkl*):

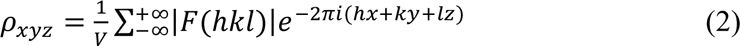

wherein *V* is the unit cell volume.

In this study, the structure factors and corresponding ED values for each complex structure were calculated using the CCTBX Python package with the pre-defined same unit cell parameters^68,69^. The binding pocket, including both protein and ligand atoms, was represented as an ED map on a cubic grid of side length 24 Å, with a spatial resolution of 0.5 Å.

To reconstruct the ligand ED from the pocket ED, we employed a VAE framework. This model consists of an encoder, which compresses the pocket ED into a one-dimensional latent variable *Z* that approximates a predefined prior distribution *p*(*Z*), and a decoder, which reconstructs the ligand ED from this latent representation. The loss function used during model training comprised two weighted components: a mean squared error term that quantified the similarity between the actual and predicted ligand ED and a Kullback-Leibler divergence term, which measured the deviation of the learned latent distribution from the true prior distribution. The loss function is formulated as:

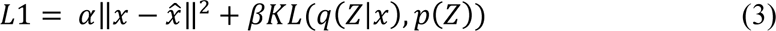

#### Initial cores placement

Our molecular generation strategy begins with fitting initial cores in the 3D space portrayed by generated ligand ED. To mitigate the computational cost associated with exhaustive searches across the entire map, we reduced the number of grid points by hierarchically applying a max-pooling operation followed by spatial clustering to identify a limited number of local density peaks. These peaks serve as candidate positions for cores placement. The cores from the library are enumerated at each identified position and systematically rotated around the XYZ axes in increments of 40°. To efficiently evaluate the compatibility of various core fragment positions and orientations with the ED map, we introduced the following Q-score score^20^:

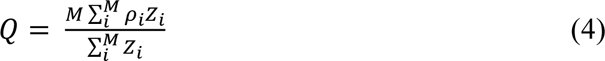

here, *Z*_*i*_ denotes to the atomic number of the atom *i*, *ρ*_*i*_ indicates the density value at the closest grid point, and *M* is the number of heavy atoms in the core. The Q-score favors conformations that exhibit strong overlap with the ED map and align with the density shape in terms of atomic number distribution.

#### Iterative fragments extension

ED space *E* is represented as a set of points {〈*c*_1_, *ρ*_1_〉, 〈*c*_2_, *ρ*_2_〉 … 〈*c*_*m*_, *ρ*_*m*_〉}, where *c* and *ρ* denote the Cartesian coordinate and the density value of grid points, respectively, and *m* is the total number of grid points. To sample a new state *s*_*i*+1_ from current state *s*_*i*_, we decomposed the process into two steps (Supplementary Fig. 1). First, the current state graph *G*_*s*_*i*__ is updated to the intermediate graph 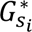 by EGNN generator part I ∅_1_, which identifies the growth site for fragment attachment. Formally:

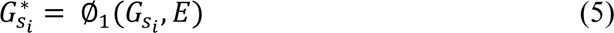

Second, given the updated intermediate state, EGNN generator part II ∅_2_ determines how to attach a new fragment graph *G*_*f*_ conditioning on *E* as follows:

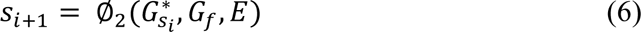

Specifically, in the first step, we encode the ED environment of atom nodes for graph embedding to assess the potential of each ligand hydrogen to serve as a growing point. For each atom node *a*, five concentric spheres with radii ranging from 1 Å to 3 Å are defined to capture the local density environment 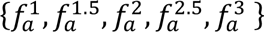. 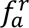 computes the number of occupied and vacant density grid points scaled by total grid points within a sphere of radius *r*, which is expressed as:

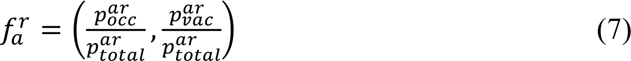

Using the local density environment as the initial atom embedding, model 1 captures atomic-level growing space information through graph convolutional layers (EGCL). Given the atom embedding *h*^*l*^ and corresponding coordinate *x*^*l*^ in the *l*^*th*^ layer, message passing is described as:

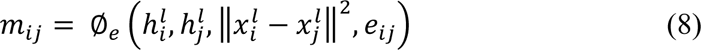

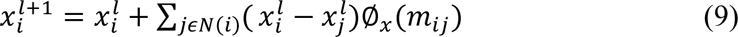

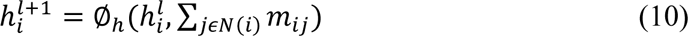

where *e*_*ij*_ is the edge feature (one-hot encoding of bond types), and ∅_*e*_, ∅_*x*_, ∅_*h*_ are multi-layer perceptrons (MLPs). Subsequent MLPs output the probabilities for hydrogen atoms, indicating whether they can provide space for new fragments. The hydrogen with the highest score is selected to attach the fragment.

In the second step, we translate the problem of determining the next state *s*_*i*+1_ into predicting the torsion angle of the newly formed bond connecting the current state *s*_*i*_and the target fragment. To this end, we adopt the normalized torsion angle *α* introduced by Ganea et al. as the model output, which considers invariances with respect to the full conformer^70^. The computation of the torsion angle *α* between bonded atoms *a* and *b* is summarized by the following equations:

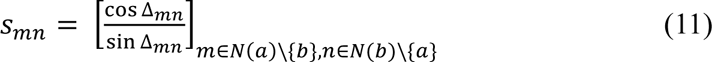

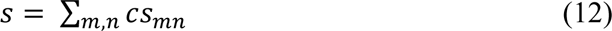

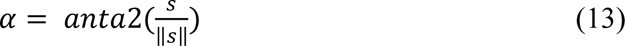

where *m* and *n* represent the local structure, the additional graph neighbors, associated with nodes *a* and *b*, respectively, and *c* refers to the real coefficients to avoid *s* to be a null vector. The EGNN generator part II incorporates three blocks, called *f*_*enc*1_, *f*_*enc*2_, *f*_*enc*3_ for separately processing different inputs. The block *f*_*enc*1_utilizes convolutional layers to map features of ED *E*, while *f*_*enc*2_ and *f*_*enc*3_ encode the intermediate state graph 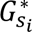 and the target fragment graph *G_f_*, respectively, extracting embeddings using EGCL layers. Please see supplementary Table 4 for details on initial feature representations of two graphs 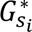 and *G_f_*. For 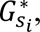 the relative atom position with the ED cube is additionally considered:

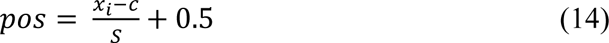

where *x*_*i*_ and *c* are the atom coordinate and center of ED, and *S* is its side length. The final torsion angle is predicted as:

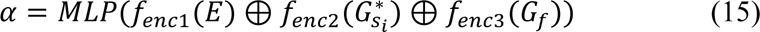

where ⊕ refers to concatenation. Each fragment in the library is fed into the model and rotated according to the predicted torsion angles. To encourage exploration of a broader chemical space, the molecules at state *s*_*i*+1_ are grouped by their originated molecules at state *s*_*i*_ and an assigned number of structurally dissimilar molecules are selected from each group based on their Q-score rankings. This strategy encourages exploration of a broader chemical space.

### Datasets construction

To construct the dataset for training the ligand ED extraction model, initially collected approximately 780,000 ligand-binding pockets from the PDB database. A subset of high-quality 49,987 pocket-ligand pairs was selected based on the following criteria:

1. Deletion of duplicated ligands appearing in multiple chains;
2. Retention of ligands with a real-space correlation coefficient greater than 0.7;
3. Exclusion of non-functional ligands and crystallization additives;
4. Selection of ligands with a molecular weight exceeding 250.

To prevent data leakage, we partitioned the dataset into training, validation, and test sets (7:1.5:1.5) such that the proteins in the test set share less than 70% sequence identity with any protein in the training or validation sets, as determined by CD-Hit (https://github.com/weizhongli/cdhit)^71^. In addition, we excluded from the training set any protein pockets that share more than 70% structural similarity with pockets in either the DUD-E or ASB-E datasets, as measured by the APOC software (https://sites.gatech.edu/cssb/apoc/)^72^.

To further build comprehensive datasets for training the models of iterative fragments extension, a subset of drug-like molecules (cLogP < 5 and 250≤MW≤500) were sourced from ZINC database for producing corresponding ED maps^5^. Each molecule was then converted into a molecular growth trajectory by decomposing it into fragments based on rotatable bonds, ensuring each fragment contained at least two atoms. Starting with the largest fragment as the initial state, the complete structure was reconstructed by iteratively attaching neighboring fragments, creating a sequence of intermediate states. Next, from these processed data, two datasets are derived, GP- and TA-datasets. The GP-dataset is designed to evaluate the potential of hydrogen atoms on an intermediate-state molecule to serve as fragment growth points. Each molecular ED map is paired with a sequence of intermediate states, labeling each hydrogen as a growth point or not. The other TA-dataset is constructed to predict the normalized torsion angles of the bonds connecting target fragments to intermediate states. The normalized torsion angle was discretized into 36 bins over the range [−*π*, *π*], enabling a classification-based prediction task. In total, the GP-dataset contains 2,986,278 samples derived from 94,519 molecules, while the TA-dataset includes 17,978,368 samples derived from 4,497,302 molecules. Both datasets were randomly split into training, validation, and test sets using a 7:1.5:1.5 ratio.

The performance of different DGMs was evaluated using two benchmark datasets: DUD-E and ASB-E. DUD-E dataset contains 102 protein-ligand co-crystal structures, primarily spanning three protein classes: GPCRs, kinases, and nuclear receptors^23^. Of these, 8 allosteric ligands, as annotated in the Allosteric Database (https://mdl.shsmu.edu.cn/ASD/), were excluded from the analysis, resulting in a final set of 94 orthosteric pockets^73^. On the other hand, ASB-E dataset was curated from ASBench (https://mdl.shsmu.edu.cn/asbench/) for validating DGMs in the more challenging scenario of discovering allosteric modulators^74^. ASBench contains 480 high-quality allosteric protein-ligand pairs, of which 296 allosteric pockets bind small endogenous ligands (e.g., ions) and were excluded due to their limited utility for structure-based allosteric modulator design. Among the remaining 184 allosteric pockets that bind exogenous ligands, 112 representative allosteric pockets were selected based on their similarity scores computed by APOC and used to benchmark the DGMs.

### Surface plasmon resonance (SPR) assay for FGFR3 inhibitors

The SPR assay was conducted using a Biacore 8K instrument (Cytiva). Human FGFR3 (residues 449-759, Biortus, USA) bearing a C-terminal Strep-tag II (BP32837-03A) was immobilized onto a CM5 sensor chip at a concentration of 50 μg/mL in 10 mM sodium acetate coupling buffer (pH 4.5) to a surface density of 18,000 Response Units (RU) via amine coupling. The surface was first activated with a 1:1 mixture of EDC/NHS (contact time: 600 s, flow rate: 10 μL/min), followed by the immobilization of FGFR3, then deactivated with 1 M ethanolamine (pH 8.5). Binding experiments were carried out at 25 °C with a running buffer comprising 10 mM Na₂HPO₄, 1.8 mM KH₂PO₄, 137 mM NaCl, 2.7 mM KCl, 0.05% Tween 20, 5% DMSO, pH 7.4. Compounds were 3-fold serially diluted using the running buffer and injected over the sensor surface at a flow rate of 30 μL/min at 25 °C with an association time of 120 s and a dissociation time of 120 s. Solvent corrections were applied using running buffers containing 4.5%–5.0% DMSO. The data were fitted with the steady-state affinity analysis in Biacore 8K evaluation software.

### Bioluminescence resonance energy transfer (BRET) assay for CDC42 inhibitors

Human CDC42 (NM_044472.2) and PAK1 (NM_001128620.2) genes were separately synthesized and cloned into a pcDNA3.1 (+) vector containing either an N-terminal Rluc or GFP tag (Sangon, Shanghai). HEK293T cells (obtained from ATCC and routinely screened for mycoplasma contamination) were cultured in Dulbecco’s modified Eagle’s medium (DMEM; Thermo Fisher Scientific, USA) supplemented with 10% fetal bovine serum (FBS). Next, the HEK293T cells were plated on 6-cm dishes at a density of 1 × 10⁶ cells/dish. After incubation for 18–24 hours, when the cells reached 60%–80% confluency, they were co-transfected with 0.4 μg of Rluc-Cdc42 and 1.6 μg of GFP-PAK1 using the ExFect Transfection Reagent (Vazyme, China) according to the manufacturer’s protocol.

The BRET assay was utilized to detect the interaction between PAK1 and GTP-bound, active form of CDC42. The transfected HEK293T cells were reseeded into white, opaque-bottom 96-well plates at a density of 10,000 cells per 100 μL per well and cultured in DMEM supplemented with 10% FBS at 37 °C and 5% CO₂ for 12 hours. The test compounds were then added to the well, and the cells were further incubated for an additional 12 hours. Afterward, the culture medium was carefully removed and replaced with Hank’s Balanced Salt Solution (HBSS; Gibco, USA) containing 5 μM Coelenterazine 400a (Maokangbio, China). Fluorescence readings were taken immediately using a Synergy Neo plate reader (BioTek, USA) with emission filters at 410 nm and 525 nm. The inhibition ratio of the test compounds was normalized to values obtained from DMSO-treated controls. The data were fitted with the standard dose-response curve in GraphPad Prism v8.0 software to determine IC_50_ values.

### Double enzyme coupled (DEC) assay for GCK activators

Human GCK (residues 1-465) was subcloned into the pET-28a vector. The recombinant protein was overexpressed in *Escherichia coli* BL21(DE3), induced with 0.5 mM isopropyl-1-thio-d-galactopyranoside at 16 °C overnight. Cells were harvested and lysed in a buffer containing 50 mM Tris-HCl pH 7.5, 300 mM NaCl and 5% glycerol. The was clarified by centrifugation at 15,000 rpm for 45 min. The resulting supernatant was applied to a Ni-NTA column (GE Healthcare, Singapore) equilibrated with the lysis buffer and then eluted with the butter containing 250 mM iminazole. The eluate was subsequently dialyzed into GCK storage buffer (25 mM HEPES, 25 mM KCl, 2 mM MgCl_2_, 1 mM DTT, pH 7.5).

The DEC assay was performed using glucose-6-phosphate dehydrogenase (G6PDH) in a final volume of 120 μl containing 25 mM HEPES pH 7.1, 25 mM KCl, 10 mM glucose, 2 mM MgCl_2_, 1 mM DTT, 1 mM NAD^+^, 0.1% BSA, 5 U/ml G6PDH, and 18.7 μg/ml GK protein. After incubation at 37°C for 30 min, the reaction was initiated by adding 1 mM ATP, and the production of NADPH was monitored kinetically at A340 nm. The activation ratio of the test compounds was normalized to values obtained from DMSO-treated controls. The data were fitted with the standard dose-response curve in GraphPad Prism v8.0 software to determine EC_50_ values.

### Crystallization and structure determination

For crystallization, the human FGFR3 (residues 449–759, Y647E, Y648E, C482A and C582S) was cloned into pET-SUMO with N-terminal 6×his tag followed by SUMO tag expressed in *Escherichia coli* BL21(DE3)-RIL. The cells were sonicated and centrifuged and the supernatant was collected. Purification was performed by sequential Ni–NTA and the soluble FGFR3 protein in supernatant was purified by affinity of Ni–NTA and cleaved by ULP1 protease in 4 °C overnight. The cleaved protein then flowed through the Ni-NTA column to remove the His-SUMO tag and followed with a gel filtration column. Final purification was achieved using a Mono Q column (GE Healthcare, Singapore), and the purified protein was concentrated to 25 mg/mL by ultrafiltration and stored at -80°C. Next, the FGFR3 protein was mixed with ACP and MgCl_2_ at a molar ratio of 1:3:15, followed by incubation for 30 minutes at room temperature. The crystal was obtained by sitting drop vapor diffusion method in 25mM HEPES pH7.5, 1.8-2.0 M LiSO_4_, 10 Mm CoCl_2_ at 20°C. Then, the crystal was soaked for 3 hours in a reservoir solution containing 1 mM test compounds before being flash-frozen in liquid nitrogen. The reservoir solution was supplemented with an additional 30% (v/v) glycerol to serve as a cryoprotectant.

The diffraction data were collected at the BL18U1 beamline of the Shanghai Synchrotron Radiation Facility (SSRF), processed with MOSFLM^75^ and merged using SCALA as implemented in the CCP4 suite^76^. The structure was solved by molecular replacement using MOLREP^77^, with the human FGFR3 structure (PDB ID: 4K33) serving as the search model. Refinement was carried out in REFMAC5^78^, applying individual isotropic restrained B-factors. Model quality was monitored using Rfree, with 5% of the reflections excluded for cross-validation. Subsequent interactive model building was performed in COOT^79^. The data collection and structure refinement statistics are summarized in Supplementary Table 5. All graphical figures of protein structure were generated using PyMOL.

## Supplementary Information

Supplementary Tables 1-5, Figs. 1-11, Methods.

